# Stiffness Sensing and Cell Motility: Durotaxis and Contact Guidance

**DOI:** 10.1101/320705

**Authors:** Jingchen Feng, Herbert Levine, Xiaoming Mao, Leonard M. Sander

## Abstract

Mechanical properties of the substrate plays a vital role in cell motility. Cells are shown to migrate up stiffness gradient (durotaxis) and along aligned fibers in the substrate (contact guidance). Here we present a simple <u>mechanical</u> model for cell migration, by placing a cell on lattice models for biopolymer gels and hydrogels. In our model cells attach to the substrate via focal adhesions (FAs). As the cells contract, forces are generated at the FAs, determining their maturation and detachment. At the same time, the cell also allowed to move and rotate to maintain force and torque balance. Our model, in which the cells only take the information of forces at the FAs, without a prior knowledge of the substrate stiffness or geometry, is able to reproduce both durotaxis and contact guidance.

## 1 Introduction

Motile eukaryotic cells can sense and react to the mechanical properties of the substrate on which they move (1–5). They do this via the focal adhesions (FA) which attach the cell to the substrate. For example, FAs on stiff substrates have been shown to be more likely to mature on flexible ones (3, 6). Cells use this information to control their motility. Durotaxis (1, 5) is an important example: it is the tendency for cells to move up a stiffness gradient. Another mechanical effect on motility is contact guidance (7, 8), the tendency for cells to move along the direction of fiber alignment. In this paper we show that both these striking effects follow naturally from a model of cell motility that properly accounts for cell mechanics and some simple properties of focal adhesions. Our model is for cell motion in two dimensions, on a substrate.

We use two different models for the substrate on which the cells move. The simplest represents the hydrogels which are often used in experiment. For this case we construct a simple two-dimensional triangular lattice. However, in real biological situations, cells move on fibrous biological gels such as collagen-I which are inhomogeneous and elastically non-linear (9, 10). For this case we use a network model that shows features such as strain-induced alignment and strain-stiffening (11–14). It is a generalization of the triangular lattice constructed by removing a fraction of the bonds (15–17).

Our model for cell motion is based on the work of Buenemann et al. (18) who considered forces and force balance in the contraction phase of Dictyostelium discoideum motility. These authors assumed a constant cell contraction rate and that the cell is connected to the substrate by adhesive bridges (intended to represent, for example, integrin) which are modeled as elastic springs. The bridges are located at the FAs. The formation of the attachments is homogeneous but detachment occurs at a spatially varying, force-dependent rate. Cell motion occurs when bridges in the back of the cell detach. Note that in this, as in all cell-motility problems, the motion is quasi-static: except for a brief interval after detachment all the forces on the cell balance. A strong prediction of this model is that the cell speed is largely independent of the value of the adhesive forces, which has been validated by a later experiment (19).

The model in Ref. (18) is essentially one-dimensional: it does not consider the reorientation of the cell during the migration process. However, in durotaxis and contact guidance cells turn in response to mechanical cues. To account for this we consider not only forces by also torques. In the course of the cell motion not only must forces (nearly) balance, but also torques. Also, in (18) the substrate is taken to be homogeneous with a constant stiffness. In this paper we generalize to the case of spatial stiffness variation.

## 2 Motility Model

In the model of Beunemann et al. (18) retraction and protrusion of actin create continuous transport of cell material to the front of the cell throughout the motility cycle. This idea is supported by the observation that cell speed is nearly a constant over the entire cycle and the motion of the cell outline is continuous sliding (20).

In the model the cell body is assumed to contract uniformly with a constant speed during the contraction phase, with duration *τ*. (Cell contraction is not hindered by viscous stress from the surrounding medium because external fluid drag is much, much smaller than the observed forces exerted on the substrate (21).) The adhesive bridges that connect the cell to the substrate form with a constant on-rate *k*_+_ and break with a force- and position-dependent off-rate *k*_−_. Their spring constants are denoted by *k_s_*.

### 2.1 Two-dimensional Mechanical Model

In this work we take the assumptions above and generalize to two dimensions by also considering torque balance. The adhesion area of a cell is modeled as an ellipse with randomly distributed sites representing FAs. The position of the center of the ellipse is called ***p**_m_*(*t*) = (*x_m_*(*t*), *y_m_*(*t*)) and the positions of the FAs with respect to the center is ***p**_i_*(*t*) = (*x_i_*(*t*), *y_i_*(*t*)). The contraction is represented by *λ* = (*A* − *A_τ_*)/*A* where *A*, *A_τ_* are the semimajor axes of the ellipse at the start and the end of the contraction. The contraction cycle is divided into 30 equal time steps *dt* (We have tried 50-time steps and the results are essentially the same). We assume the contraction only occurs along the long axis. The orientation of the cell at time *t* is called *θ*(*t*). Then, from simple geometry, the contraction dynamics of the node *i* is:

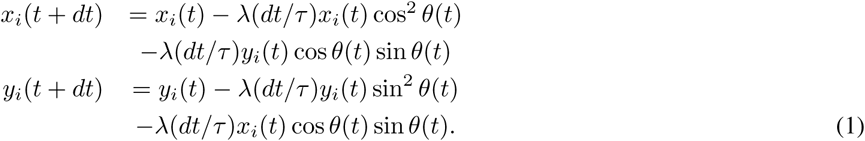

At the beginning of the contraction phase, FAs are formed at each network node within the adhesion area with probability *k*_+_. The force on a single FA is given by:

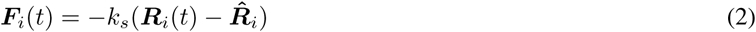

Here, ***R****_i_* = ***p****_i_* + ***p****_m_* is the position of FA *i* and 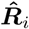 the position of the network node. The total energy of the springs at the FAs at time *t* + *dt* is:

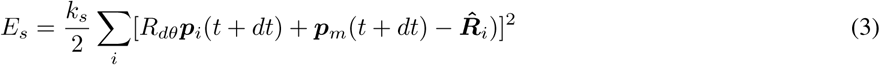

Here *R_dθ_* is the 2D rotation matrix through *dθ*. The derivative of *E_s_* with respect to *θ* is the net torque on the cell, and the derivative with respect to ***p**_m_*(*t* + *dt*) is the net force on the cell. The cell center ***p**_m_* and cell orientation *θ* are allowed to move to ensure zero net force and torque. At the same time, the network nodes 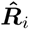 are also allowed to move, minimizing the total energy of the cell and the network elasticity, as we discuss below.

Once the cell contracts, the traction forces on FAs will build up and a number of FAs will detach. In order to account for cell polarization, we need the attachments to be weaker (i.e. have a larger off-rate) at the back of the cell than in front. We encode this as follows:

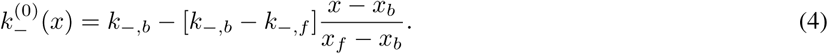

Here *x_f_*/*b* are the front/back of the cell at the start of the contraction cycle and *k*_−,_*_f/b_* are parameters with the constraint *k_b_* > *k_f_*.

The force dependence of the off-rate is modeled by Bell’s law where the off-rate exponentially grows with the stretching (18, 22):

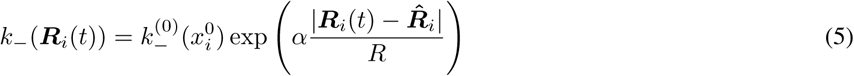

**Figure 1:**
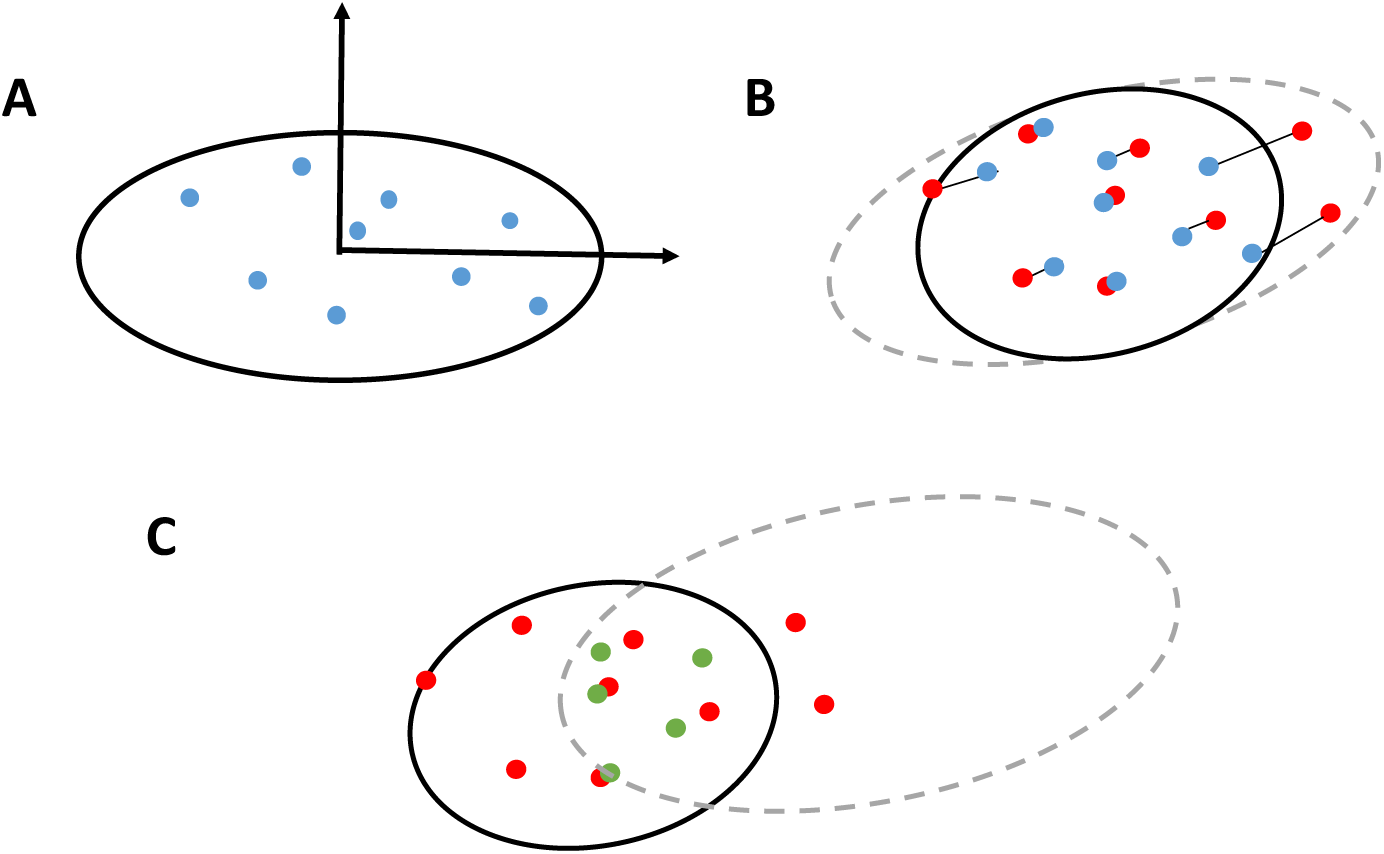
Representation of cell-substrate adhesion during the contraction cycle. (a) The start of the contraction cycle. FAs are blue circles. The FAs form randomly at network nodes. (The network structure is not explicitly shown here.) (b) During the motility cycle, the cell contracts uniformly at a constant speed. The position of the attached network nodes is shown as red circles. The current position of FAs are blue circles. The contraction causes deformation of the network and rotation and shift of the cell. (c) The end of the contraction cycle: remaining FAs are shown in green. At the start of a new motility cycle, the cell outline is shifted such that its back coincides with last remaining adhesion site as indicated by the dashed ellipse.

Here 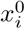 is the initial position of the FA along the major axes of the ellipse. The dimensionless parameter *α* measures the strength of the bond: see (18). At the start of a new motility cycle, the cell outline is shifted such that its back coincides with last remaining adhesion site.

The network model that we use for biopolymer gels is known to reproduce many important features of fibrous gels such as strain-induced alignment and strain-stiffening (11–14). The network is built on a diluted triangular lattice as shown in Figure (2). Each bond in the lattice is present with a probability *p*. The probability *p* satisfies *pZ* = 〈*z*〉, where *Z* is the coordination number of the undiluted lattice (*Z* = 6 for triangular lattice), and *z* is the average connectivity of fibrous network. In experiment 〈*z*〉 ≈ 3.4 (23). Therefore we study *p* in the range [0.5,0.65], We make contact with the mechanics of physical biopolymer gels by identifying the lattice sites as cross-linking points and bonds as fibrils between crosslinks. For simpler substrates such as hydrogels we simply put *p* = 1.

**Figure 2:**
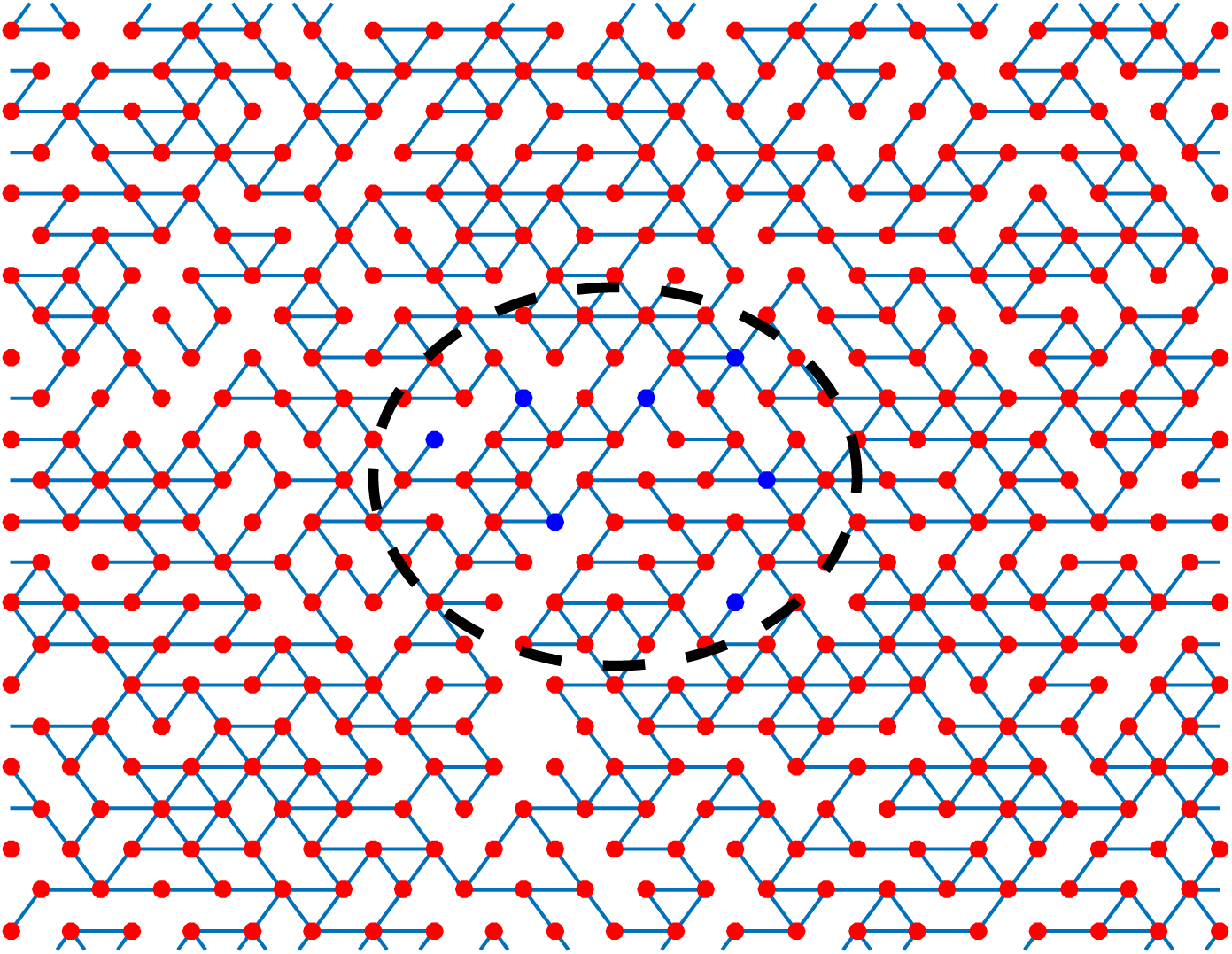
The network model. The red circles are network nodes (crosslinking points) and the blue bonds are the fibrils between crosslinks. In the beginning of the contraction cycle, FAs (blue circles) have a probability to form on the top of each nodes within the adhesion area (dashed ellipse). Periodic boundary condition is applied to the network model in all the simulations

The elastic energy of the network is:

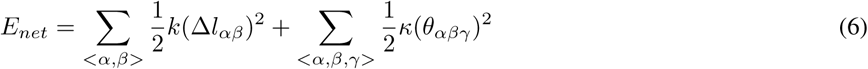

The sum < *α*, *β* > runs over bonds and < *α*, *β*, *γ* > runs over pairs of bonds that are co-linear and share lattice site *j*. The length change of the bond is 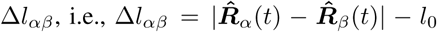 where *l*_0_ is the rest length of the bond. The spring constant of the bonds is *k* and *κ* is the bending stiffness. We always take 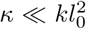 for fibrous gels, in agreement with experiment.

Cell contraction results in the displacement of the FAs, which leads to a nonzero net force and torque on the cell. To restore mechanical equilibrium we minimize the total energy of the system *E_total_* = *E_s_* + *E_net_* with respect to the position and orientation of the cell. That is, the cell (ellipse) is allowed to shift and rotate, and the network is allowed to deform.

### 2.2 Model with FA maturation

The direct two-dimensional generalization of (18), described above, captures the fact that the FAs detach more efficiently on a stiffer substrate because the forces at the FAs are stronger, and as a result, cells move faster on a stiff substrate. However, it is not sufficient for our purposes. On some substrates, as we will see below, it shows durotaxis in a natural way, but in other cases cells migrate down the stiffness gradient, which is never observed. However, this cannot be right: durotaxis, for example, has a biological function (5) which needs to be robust in many biological environments.

Note that there is a well-known property of FAs, namely that they are more likely to mature on stiff substrates (3, 6). For example, a recent experiment (6) shows that the local matrix microenvironment regulates the adhesion lifetime. It is positively correlated with the stiffness of Extra-Cellular Matrix. By inserting this effect into our model we make it robust for all the substrates we have examined.

Thus we assume that the maturation of the focal adhesions is dependent on the local stiffness of the substrate. In our study, the cell probes the local stiffness by contracting and the maturation of FAs depends on the traction forces between FAs and the substrate. Our model with FA maturation does show positive durotaxis as observed in experiments. It also shows that the orientation of the cell is along fiber alignments, which is consistent with experimental observations (7, 8) of contact guidance.

In detail, instead of assuming all FAs survive after formation, we assume that nascent FAs are formed at each network node within the adhesion area with the probability *k*_+_. At the end of the first time step *dt*, we determine if the FA will mature or perish based on the loading force on the FA. The maturation probability is taken to be:

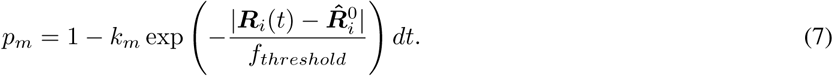

For the rest of the contraction phase, mature FAs can detach from substrate at the end of each time increment, as we do in the basic model.

### 2.3 Stiffness gradients and fiber alignment

We first simulate the case where cells are placed on the top of a simple substrate such as hydrogel as in many *in vitro* experiments. We represent hydrogel as a network with *p* = 1 and introduce a stiffness gradient by varying the spring constant k in space. To create a sharp jump in stiffness gradient, we take the upper half of the network to have *k_upper_* and the lower half of the network *k_lower_*: Figure (3a).

**Figure 3:**
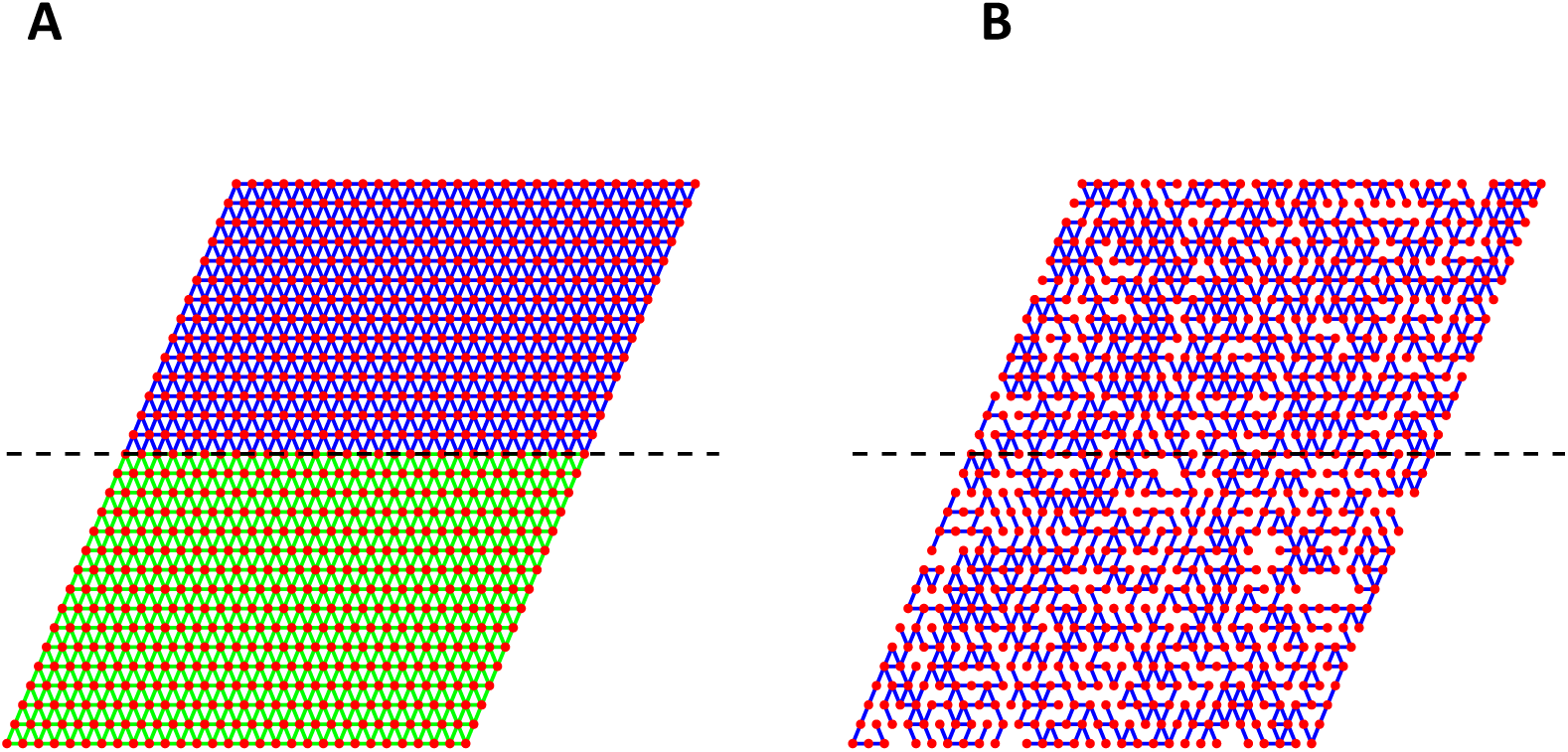
Setting a stiffness gradient in the network model. (a) Hydrogel case (p=1). *k_upper_* = 1.00 (blue) and *k_lower_* = 0.001 (green). (b) Fibrous network case. *p_upper_* = 0.65 and *p_lower_* = 0.50 The dashed line shows the location of the interface.

In contrast, for a biopolymer gel such as collagen, cells are placed on the top of the diluted network. We introduce a stiffness gradient by keeping *k* constant but spatially varying *p* – in effect we are modeling density variations in the gel. We create the sharp jump in by putting *p* = *p_upper_* in the upper *p_lower_* in the lower part: Figure (3b).

For the contact guidance case we use the fact that gels have strain-induced alignment (11). In the simulations we introduce fiber alignment by simply stretching the network: Figure (4).

**Figure 4:**
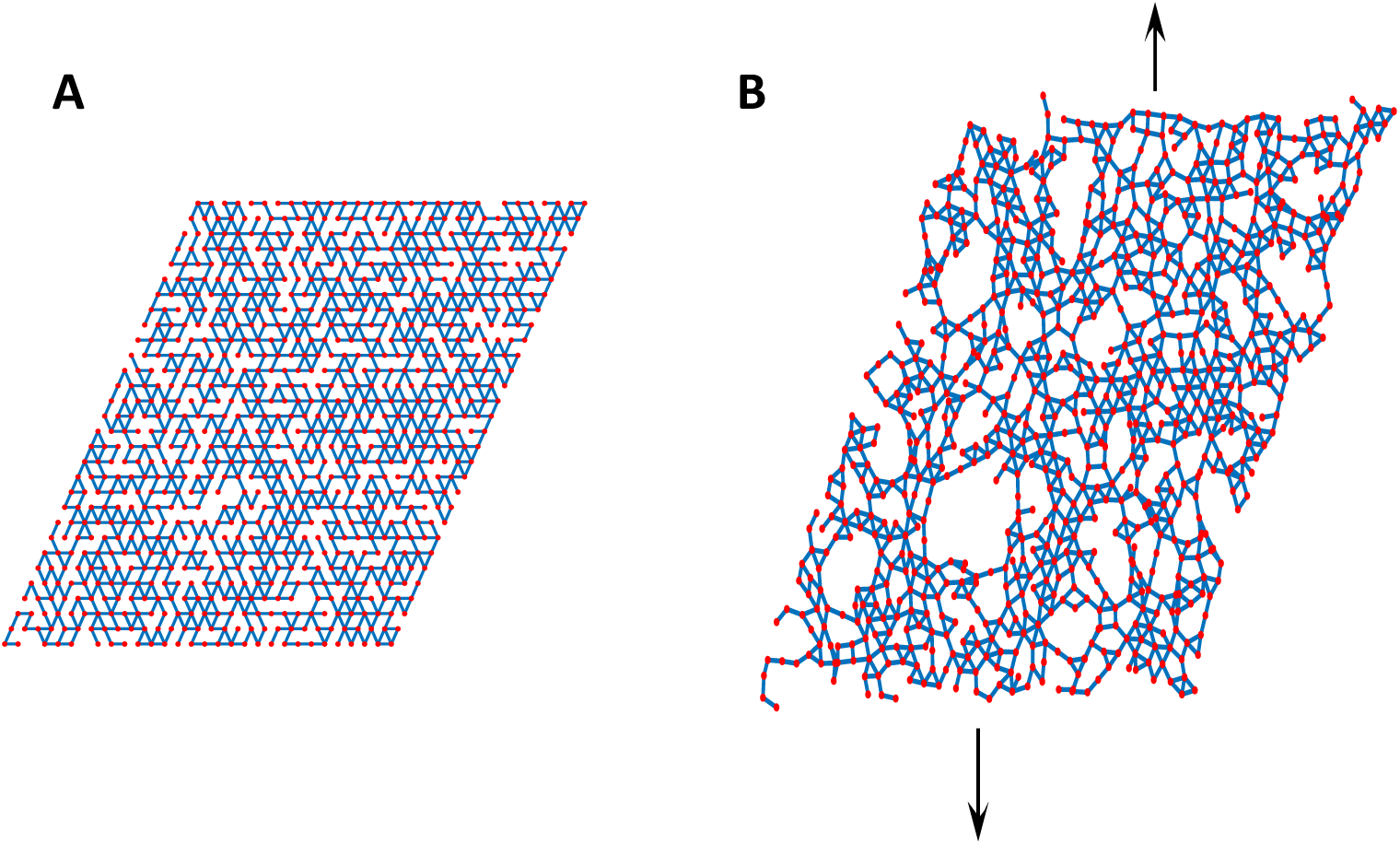
Fiber alignment. (a) The rest state of the network. (b) Fibers are aligned along the stretching direction by external deformation. Here *p* = 0.60, *κ/k* = 0.001, stretching strain *γ* = 0.40.

## 3 Results

### 3.1 Durotaxis

We now turn to simulations which show durotaxis. We first test the hydrogel case (*p* = 1) where sharp jump in stiffness is created by varying spring constant *k* as in Figure (3). We start the cell on the interface with orientation parallel to the interface. The simulation lasts for 15 cell cycles. At the end of the simulation, we record the location and the orientation of the cell.

We find that in the basic model (without FA maturation) the cell will move towards decreasing stiffness, which is the opposite of durotaxis: see Figure 5(a). This is easy to understand: suppose a cell has four FAs at top left, top right, bottom left and bottom right respectively. The FAs receive traction forces from the network due to cell contraction, and the cell is in torque balance. At a certain point, the top left FA will break first, since the cell has a larger detachment rate at the rear and the stiff region causes a stronger stretching on FAs. This breaking event will further causes torque imbalance. The cell needs to rotate clockwise to reach new torque balance (Figure 5(b)). FA maturation resolves this problem sicne more FAs mature in the stiff region, and the cell with can correctly sense and move towards the stiffer region (Figure 5(c)). In the following, we will use the full model with FA maturation unless otherwise stated.

**Figure 5:**
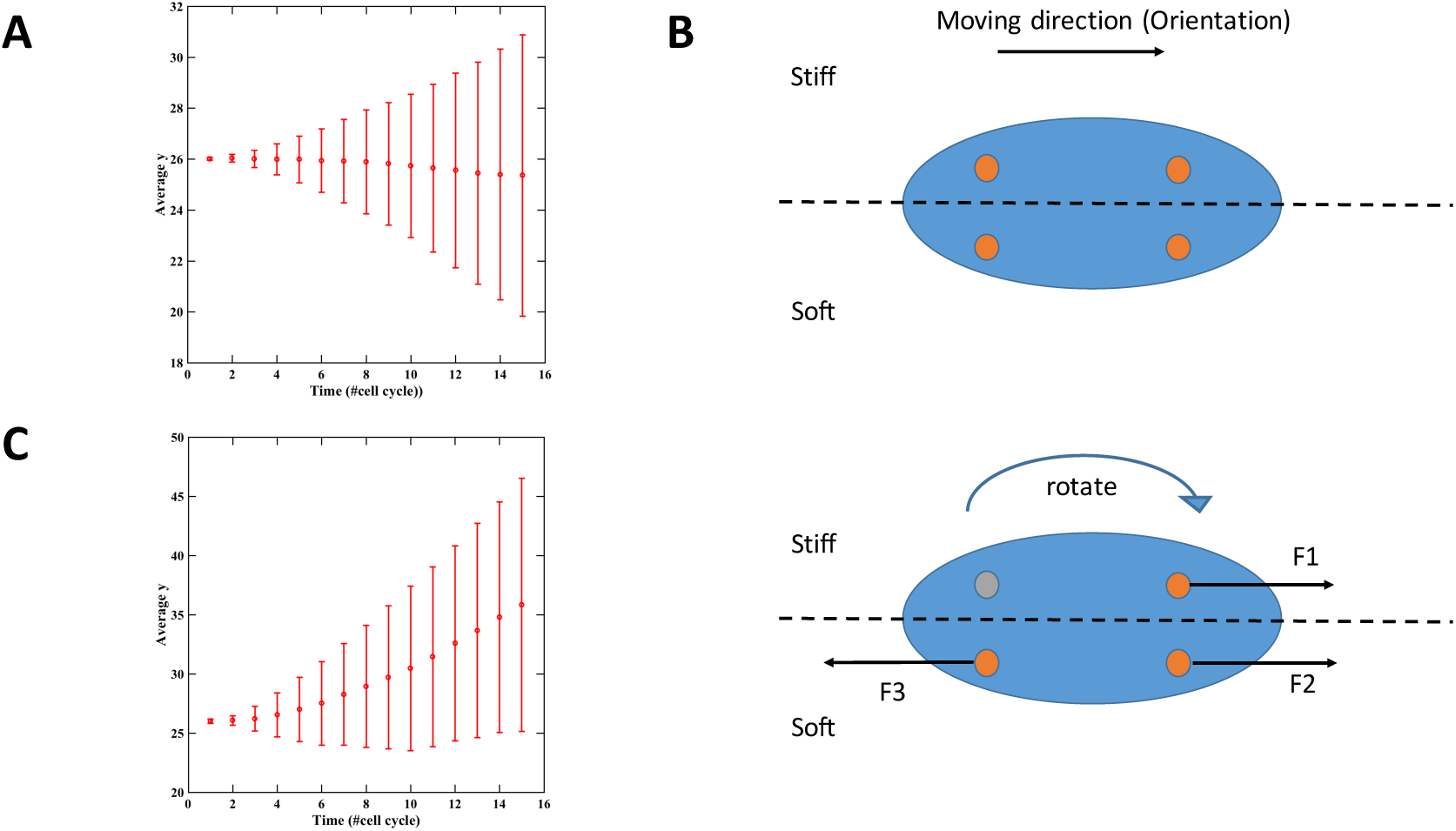
Stiffness gradient by varying *k*. The average *y* coordinate as a function of time (a) The cell model without FA maturation shows negative durotaxis. (c) With FA maturation the cell moves towards the stiff (top) region. (b) Cell orientation change in the model without FA maturation.

Next we test the biopolymer network case. Figure (6) is an example of a cell (with FA maturation) moving on a substrate with a spatially varying stiffness by varying *p*. Initially, the cell center is on the interface: Figure 6a. After two cell cycles, the cell moves rightwards but does not steer to the stiff region yet Figure 6b. After four cell cycles, the cell reorients towards the stiff region: Figure 6c.

**Figure 6:**
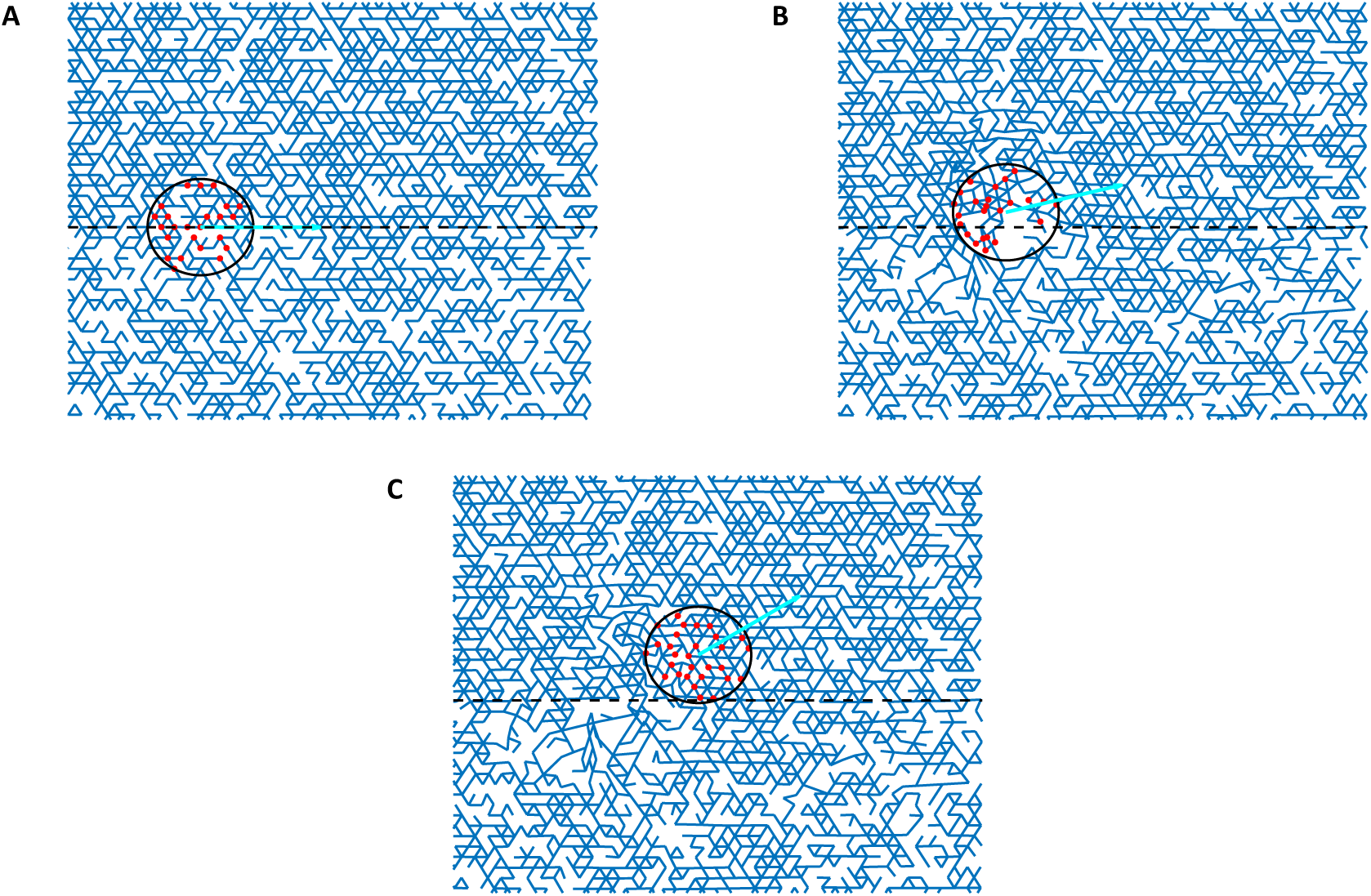
An example of a cell (with FA maturation) migrating on a substrate with a spatially varying stiffness. (a) initial state; (b): third; (c); fifth cell cycle. The red dots represent focal adhesions. Dashed line shows the interface between the stiff substrate (top) and soft substrate (bottom). *p_upper_* = 0.65, *p_lower_* = 0.50, *κ/k* = 0.001. Arrows indicate the orientation of the cell. Note that in this example the cell is circular (radius equals 4 lattice spacing). The actual shape of the network is the same as in Figure 4. The right side of the network is pieced together with left side in the figures above, since the periodic boundary condition is applied.

In Figure (7) we repeat the simulation 500 times and measure the average coordinate of the cell, 〈*y*〉, and the distribution of cell orientations at the end of the simulation. It’s clear that 〈*y*〉 and 〈*θ*〉 is positively correlated with the stiffness gradient. The cell’s ability is highly sensitive to stiffness gradients. For example, for *p_upper_* = 0.60 and *p_upper_* = 0.55, the geometry of the two sides of the network looks essentially the same (see Supplementary Materials). Nevertheless, the cell shows a strong preference to move up (Figure 7e). Since we use a large number of FAs the model integrates the stiffness information over the whole cell adhesion area and averages out the randomness. As a result, the cell shows a consistent tendency to move towards the stiff region.

**Figure 7:**
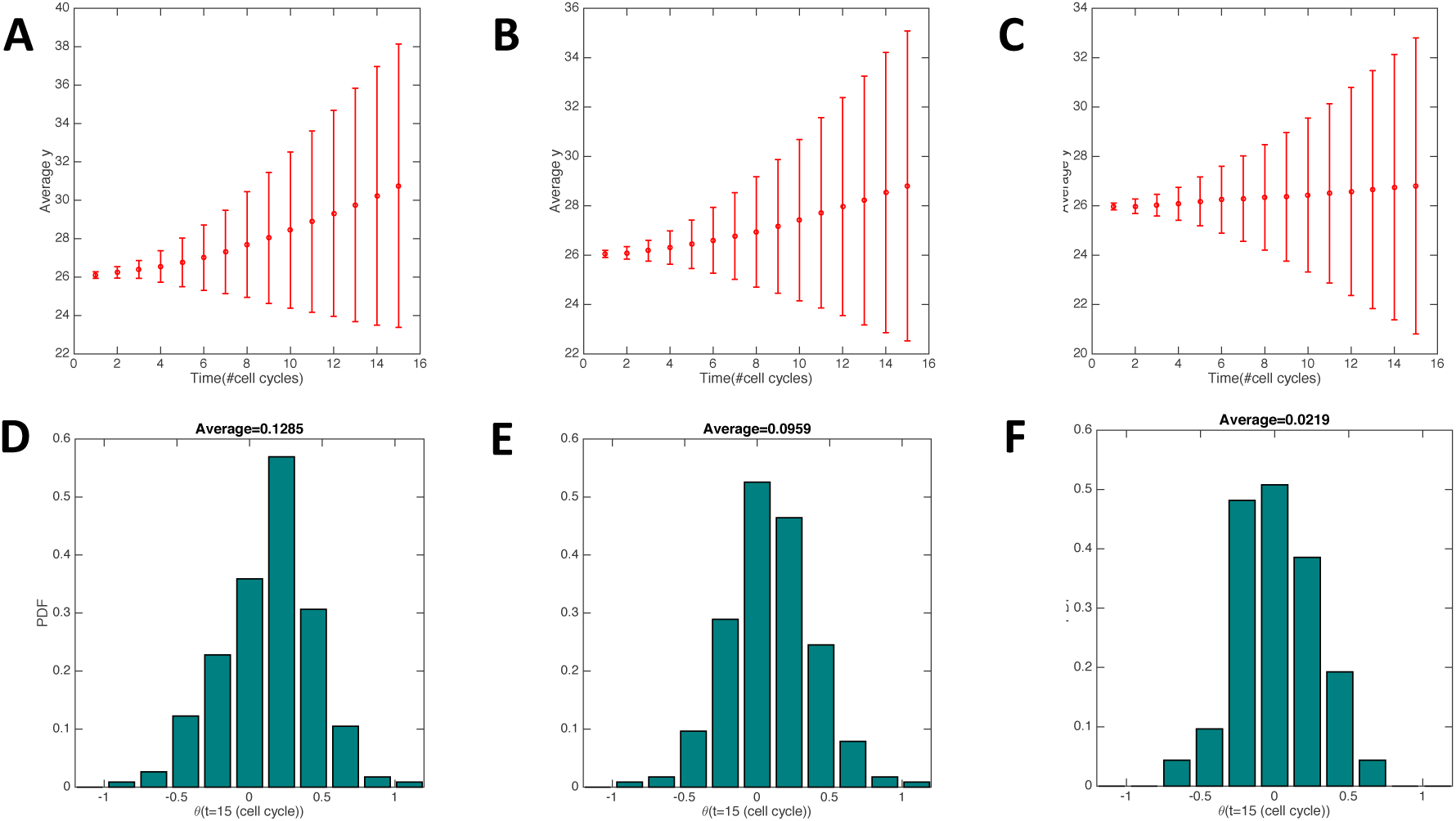
Stiffness gradients guide cell migration. (a)-(c) Average *y* coordinate of the cells versus time. (d)-(f) Distribution of cell orientations. Parameters: *κ/k* = 0.001, (a) and (d) *p_upper_* = 0.65, *p_lower_* = 0.50. (b) and (e) *p_upper_* = 0.60, *p_lower_* = 0.55. (c) and (f) *p_upper_* = 0.58, *p_lower_* = 0.57

Oddly, in this case we get essentially the same results for the model without FA maturation because a network with *p* < 2/3 (below the central-force isostatic point) (16) is highly deformable. The cell reorientation is induced by the local network remodeling. Therefore, FA maturation does not play an important role.

### 3.2 Contact guidance

We now turn to contact guidance, namely the tendency of cells to follow fiber orientation (7, 8). We first stretch the network in the vertical direction with strain *ϵ* to induce fiber alignment. Then a cell is put on the substrate with a uniformly distributed orientation *θ*(*t* = 0). The simulation lasts for 50 cell cycles and the orientation of the cell is recorded at the end of the simulation. The simulation is repeated 500 times. To quantify the alignment, we calculate the nematic order parameter *N* for both cells and fibers in each samples. *N_cell/fiber_* = 〈cos(2*θ*)〉, where *θ* is the direction of the cell/fiber orientation measured from the stretching direction.

Figure 8a shows that both fiber alignment *N_fiber_* and cell orientation *N_cell_* are positively correlated with the stretching strain *ϵ*, confirming both strain-induced alignment and contact guidance.

**Figure 8:**
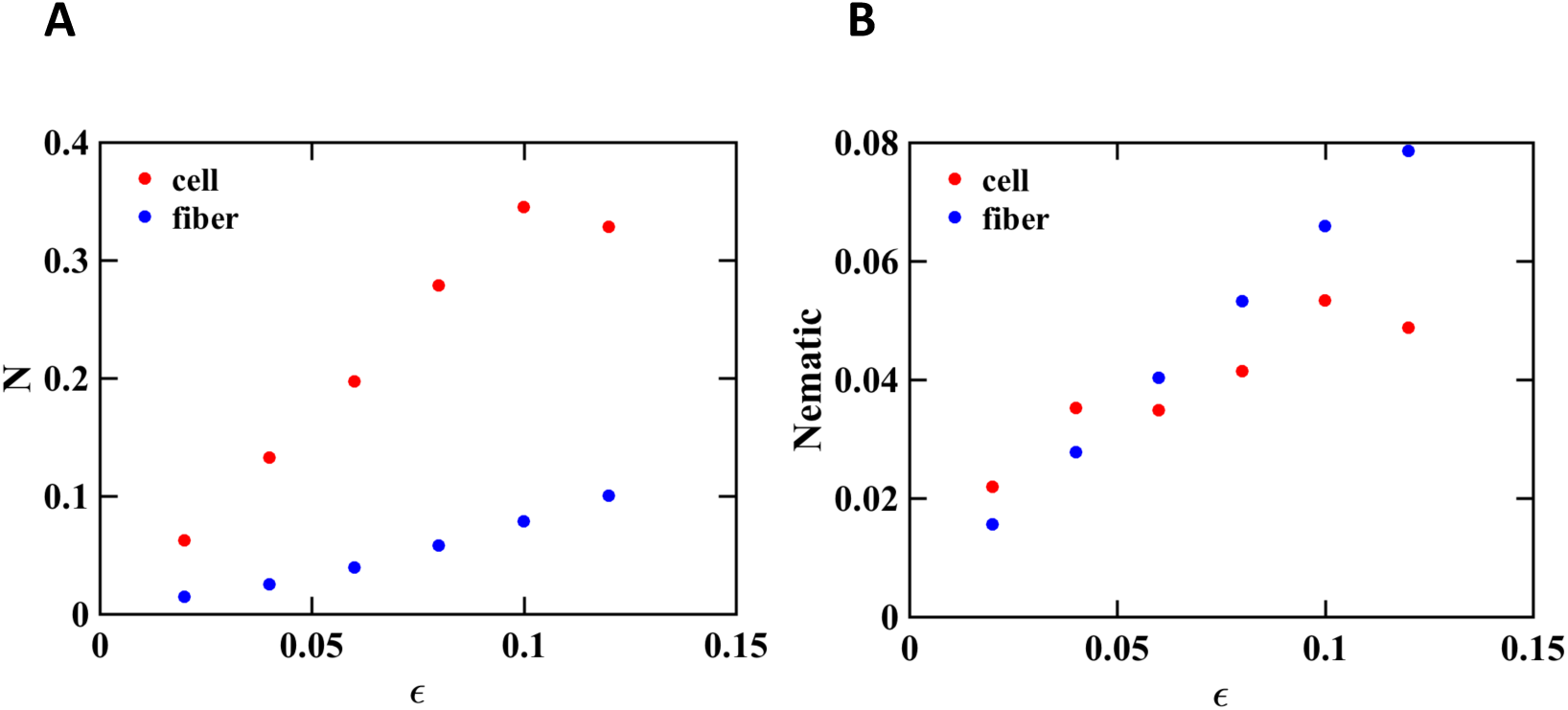
Nematic order parameter *N* for cells and fibers. *p* = 0.60 and (a) *κ/k* = 0.001 (b) *κ/k* = 1. *N_cell_* is calculated over 500 cell samples, and *N_fiber_* is calculated over all fibers in each sample.

Note that external strain induces both fiber alignment and stiffness anisotropy. To figure out which is important in contact guidance in our model, we test a slightly different case where the bending stiffness *κ* = 1 (Figure 8b). Interestingly, the strain-induced fiber alignment stays largely the same in this case, while *N_cell_* is strongly suppressed. This observation suggests that the network geometry alone is not enough for contact guidance, because in both *κ* = 1 and *κ* = 0.001 cases we observe comparable fiber alignment strength, but the cell orientation is quite different. A plausible explanation here is that the cell senses the stiffness of its environment by deforming it. When the network is too stiff to deform, the cell loses its ability to sense stiffness anisotropy, resulting in the weakening of the contact guidance. In cases where the geometry is very anisotropic, it is possible that contact guidance could occur from geometry alone, as well (24).

The distribution of the cell orientation (Figure 9(a)-(c)) shows two peaks in the probability density function, illustrating cells’ preference in moving along the fiber alignment direction. Such distribution can also be understood with a Fokker-Planck equation (See Supplementary Materials). The dashed lines in Figure 9(a)-(c) are the fitting results of the Fokker-Planck approach. This alternative Fokker-Planck approach also gives consistent value of the initial slope of the curve *N_cell_* vs. *N_fiber_* to the simulations. We leave the discussion of Fokker-Planck approach in the Supplementary Materials because Fokker-Planck approach does not consider the mechanical aspect of cell migration, which is the highlight of this paper.

**Figure 9:**
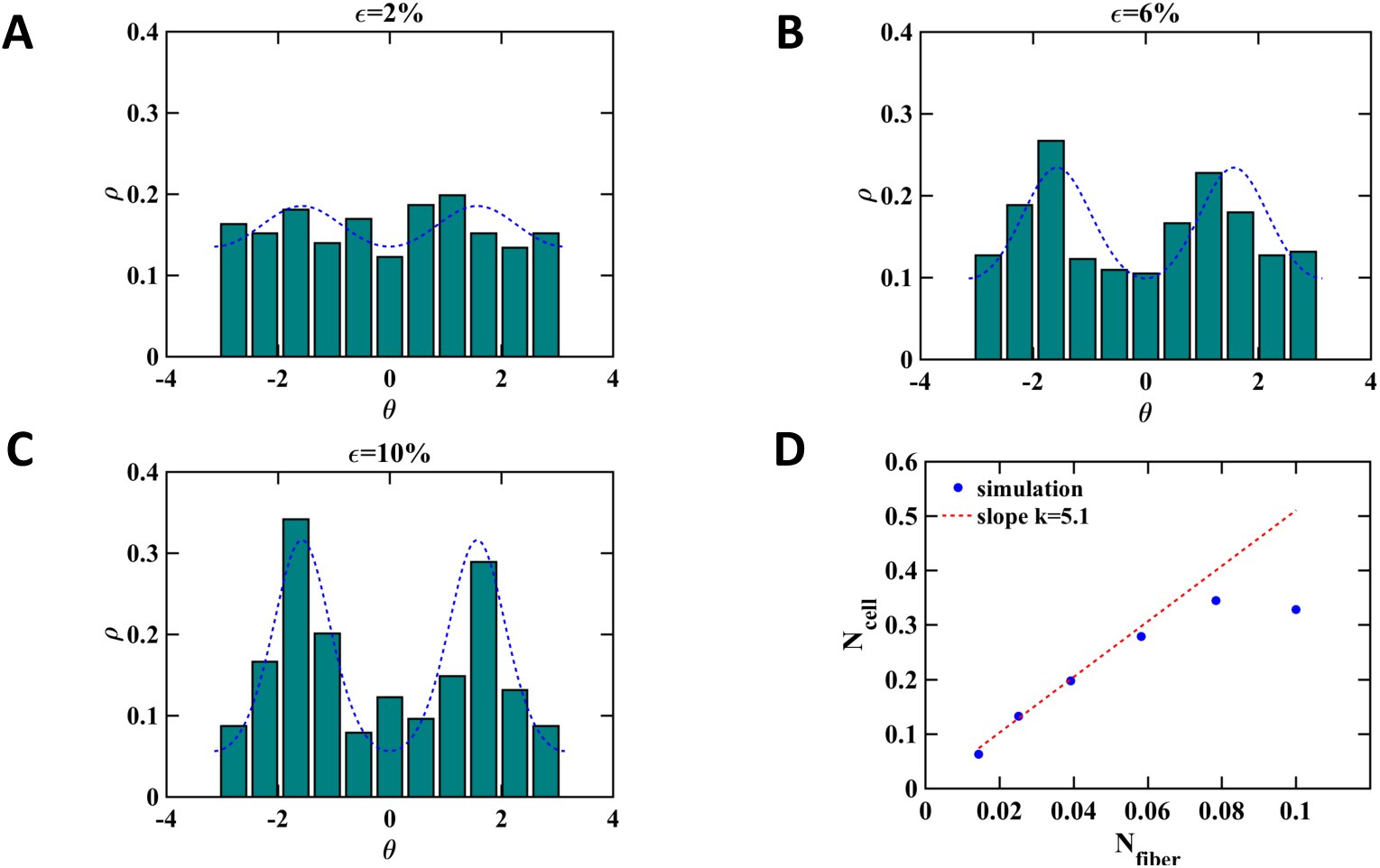
(a)-(c) cell orientation distribution after 50 cell cycles. The blue line represents the results of Fokker-Planck equation with 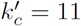 (See Supplementary Materials). Stretching strain (a) *ϵ* = 2% (b) *ϵ* = 6% (c) *ϵ* = 10%. (d) nematic order of cell orientation versus nematic order of fiber alignment. The dashed line is a linear function with a slope of 5.1.

## 4 Discussion

In this paper we present a simple mechanical model basing on the work of Buenemann et al. (18). By generalizing the model to two-dimension, the model considers torque balance of the cell in addition to force balance. This naturally allows the cell in our model to re-orient during the migration. Consistent with experimental observations, our model exhibits the reorientation is influenced by both fiber alignment and stiffness gradient. Our model also shows that FA formation, maturation and detachment play a critical role in determining the reorientation of the cell.

Most early experiments reporting durotaxis were conducted on hydrogels instead of biopolymer gels (1, 25). However, biopolymers constitutes a major part of tissues. It is important to consider the fibrous nature of tissues (26). A main difference between hydrogels and biopolymer gels is that biopolymer gels usually has strong mechanical anistropy (27). To address both scenarios, we model hydrogels and biopolymer gels as lattice-based models with different bond occupation probability. We find that FA maturation is indispensable for the case of durotaxis on hydrogels. Actually, we show with a simple model that the cell always prefers to move along the descending direction of stiffness in the absence of FA maturation in this case. In contrast, we observe that FA maturation is negligible in the case of biopolymer gels. One possible explanation is that biopolymer gels are highly deformable so that the traction forces of the cell cause local remodeling of the gels. The remodeling may be responsible to durotaxis without FA maturations.

As for contact guidance, an interesting question is whether the geometrical anisotropy or the mechanical anisotropy is the fundamental cause. To answer this question, we first want to emphasis that mechanical anisotropy is inseparable from geometrical anisotropy. To give a simple example, let’s consider a gel with all fibers aligning in the horizontal direction. In this situation, we can easily see that the gel is easier to deform in the vertical direction than the horizontal direction, namely mechanical anisotropy originates from geometrical anisotropy. In our simulation, we vary the bending stiffness *κ* of the network to generate networks with similar geometry but different mechanical property (Figure 8b). Interestingly, we observe that the contact guidance of the cell disappears in the large *κ* limit, indicating that the geometry has to act through mechanical response and mechanical anisotropy is the fundament cause of contact guidance.

The aim of this paper is to show that a simple mechanical model is capable of explaining both durotaxis and contact guidance. In reality, cell migration is complicated by many chemical reactions and biological processes. For instance, mechanical signaling can regulate cellular behaviors via signaling pathways(28, 29). Some cells can secret Matrix Metalloproteinases (MMPs) which can remodel extracellular matrix proteins (30). In this work we simplify the problem by considering the mechanical aspects of cell migration only. Our model can serve as a framework in cell migration modeling. In future work, chemical reactions and biological processes can be incorporated into our model to study more complex scenarios.

Simulating cell and substrates interactions is computationally challenging. As a starting point, we study population-level phenomena, contact guidance and durotaxis, by repeating single cell simulations. The defect of this approach is that we neglect the cell-cell interactions completely. Therefore our model can only be used to understand cell migration in low cell density situations. We plan to conduct multi-cell simulations and study the role of cell-cell interactions in cell migrations in the near future.

## SUPPLEMENTARY MATERIAL

An online supplement to this article can be found by visiting BJ Online at http://www.biophysj.org.

## Acknowledgements

J.F. and H.L. are supported by the National Science Foundation Center for Theoretical Biological Physics (Grant PHY-1427654). X.M. is supported by the National Science Foundation Grant NSF-DMR-1609051. H.L. is also supported in part by the Cancer Prevention and Research Institute of Texas Scholar Program of the State of Texas at Rice University.

